# A self-adjusting, progressive shock strength procedure to investigate resistance to punishment: characterization in male and female rats

**DOI:** 10.1101/2022.03.12.481595

**Authors:** Stevenson Desmercieres, Virginie Lardeux, Jean-Emmanuel Longueville, Myriam Hanna, Leigh V. Panlilio, Nathalie Thiriet, Marcello Solinas

**Affiliations:** Université de Poitiers, INSERM, U-1084, Laboratoire des Neurosciences Expérimentales et Cliniques, Poitiers, France; Real-world Assessment, Prediction, and Treatment Unit, Translational Addiction Medicine Branch, Intramural Research Program, National Institute on Drug Abuse, Baltimore, MD, USA

**Author notes:** **Corresponding author:** Dr Marcello Solinas, Université de Poitiers, INSERM U-1084, Laboratoire des Neurosciences Expérimentales et Cliniques 1, rue Georges Bonnet, TSA 51106, 86073 Poitiers Cedex 9, France.

**Keywords:** punishment, reward, compulsivity, punishment, addiction, motivation

## Abstract

Indifference to harmful consequences is one of the main characteristics of compulsive behaviors and addiction. Animal models that provide a rapid and effective measure of resistance to punishment could be critical for the investigation of mechanisms underlying these maladaptive behaviors. Here, analogous to the progressive ratio (PR) procedure widely used to evaluate appetitive motivation as the response requirement is increased, we developed a self-adjusting, progressive shock strength (PSS) procedure. The PSS provides, within a single session, a break point that quantifies the propensity to work for a reward in spite of receiving electric footshock that progressively increases in duration. In both male and female rats, the PSS break point was sensitive to 1) hunger; and 2) changes in the qualitative, but not quantitative, incentive value of the reward. In systematic comparisons between PSS and PR procedures in the same rats, we found that both measures are sensitive to manipulations of motivational states, but they are not intercorrelated, suggesting that they measure overlapping but partially distinct processes. Importantly, the PSS procedure represents a refinement in the 3Rs principles of animal research because animals can control the strength of shock that they are willing to tolerate. This self-adjusting PSS procedure may represent a useful tool to investigate mechanisms underlying maladaptive behavior that persists in certain individuals despite harmful consequences.

## 1 Introduction

Addictive and compulsive disorders are characterized by the urge to execute behaviors even when they cause pain, suffering, or health problems (Figee et al., 2016). In rodents, compulsive behaviors are often mimicked by resistance to punishment in the context of food or drug-taking behavior. The most commonly used punishment procedures use a fixed strength of foot-shock which allows identifying animals that are sensitive or insensitive to shock (Belin et al., 2011; Deroche-Gamonet et al., 2004; Domi et al., 2021; Marchant et al., 2018; Pascoli et al., 2015; Pelloux et al., 2007). However, using a fixed level of shock imposes a dichotomy based on an arbitrary threshold; it labels each animal as either sensitive or resistant, without taking into account the fact that sensitivity is a continuous variable. That is, every individual has some level of shock where it will switch from responding to not responding, and this threshold may vary between individuals and over time within the same individual. A more nuanced measure of individual resistance to punishment can be obtained with procedures where the strength of foot shock is increased over several sessions (Durand et al., 2021; Krasnova et al., 2014; Panlilio et al., 2003) or in blocks within a single session (Bentzley et al., 2014; Datta et al., 2018) until animals decrease and eventually stop responding. However, even in these procedures the levels of shock are imposed by the experimenter in a fixed (increasing) order and animals have little control of the strength of shock that they are willing to accept to obtain rewards. Experimenter-imposed procedures involve the risk of delivering shocks that are substantially higher than what the individual would be willing to tolerate, which can lead to persistent inhibition of reward seeking (Durand et al., 2021). Indeed, punishment with high levels of shock on first encounter can dramatically suppress operant responding whereas punishment with lower increasing levels of shocks has more gradual effects on responding (Miller, 1960). Procedures that adjust the level of shock to individual behavior may allow a more precise measure of resistance to punishment and be more in accordance with ethical standards because punishment is titrated within a range close to the individual’s threshold, only exceeding it infrequently and by a small amount. In addition, they would resemble procedures commonly used to investigate the effects of punishment in humans, in which shock levels are calibrated individually (Apergis-Schoute et al., 2017; Kanen et al., 2021; Kim and Anderson, 2020).

Progressive ratio (PR) schedules (Hodos, 1961) are considered the gold-standard for the investigation of incentive motivation and the strength of rewards. In PR schedules, the number of responses required to obtain a reward increases at each step until the animal stops responding, and the final step, or break point, is considered a direct measure of individual motivation. One of the advantages of PR schedules is that break points can be obtained within a single session. PR schedules for food rewards are sensitive to the motivation of the individual in terms of hunger (Hodos, 1961; Solinas and Goldberg, 2005) and of incentive value of the reward (Hodos, 1961). To incorporate the advantages of PR procedures, we designed a self-adjusting progressive shock strength (PSS) procedure in which the strength of shock is adjusted within a single session according to the animal’s behavior. In this procedure each animal can titrate its own “acceptable” level of punishment to obtain food rewards.

The psychophysical experience of shock and its punishing effects depend on the intensity (current) and the duration of footshocks (Church et al., 1967; Leander, 1973). These two variables can be manipulated interchangeably (Leander, 1973) determining the electrical charge delivered, which can be considered a measure of the strength or magnitude of shock. Changing the intensity of a shock remotely within a session is tricky and requires specialized equipment. In contrast, changing the duration of footshocks can be easily and precisely implemented with conventional shockers by simply altering a parameter in the software program used to control the operant procedure. Thus, in our procedure, the strength of shock was determined by using a fixed shock current (0.45 milliamperes, mA) and manipulating the shock duration (from 0 to 0.71 sec), leading to electrical charges ranging from 0 to 0.3195 milliCoulomb (mC).

Both resistance to punishment and PR motivation can be viewed as measures of the costs that an individual is willing to accept to obtain a reward. Interestingly, Hodos developed his PR procedure to quantify motivation in order to overcome some technical problems that had led to excessive variability in the results obtained in older progressive shock procedure (Hodos, 1961). Although motivation PR and resistance to punishment are often associated (Hogarth, 2020), the relationship between these measures is not straightforward. Indeed, several studies have found that resistance to punishment and PR responding do not always correlate (Datta et al., 2018; Pascoli et al., 2015; Pelloux et al., 2007), suggesting that the willingness to exert an effort or the willingness to tolerate a negative outcome to obtain a reward represent two somewhat different aspects of motivation. To gain insights into these aspects of motivation, we systematically compared the effects of manipulations of motivation on the PSS and PR procedures. We tested whether the punishment break point is sensitive to manipulations of motivational states such as hunger, reward size and reward quality, and we evaluated the effects of these manipulation on progressive ratio responding.

## 2 Methods

### 2.1 Subjects and housing conditions

Sprague-Dawley rats (48 males and 24 females) aged 8-9 weeks (Janvier Labs, France) were used in this study because rats’ behavioral repertoire makes them a species of choice for the investigation of addiction-related processes (Parker et al., 2014). The experiments were performed in 3 separate cohorts of animals: 1) 24 males to establish the procedure; 2) 24 males to replicate and assess reliability of the procedure and 3) 24 females to assess whether the results depend on sex. It should be noted that the female cohort and the 2 cohorts of males were all run separately. Results from the two male groups were basically superimposed qualitatively and quantitatively suggesting that the conducting the experiments in separate cohorts had minimal effects on the results. Data from the two cohorts of males were combined for analysis and presentation here. Rats were housed in pairs with two wooden sticks for enrichment, in IVC cages (Techniplast, Sealsafe Plus GR900), in a temperature- and humidity-controlled room kept under reversed light-dark cycle conditions (12 hours light-dark cycle, lights off at 7 AM). All experiments were conducted during the dark phase and in accordance with European Union directives (2010/63/EU) for the care of laboratory animals and approved by the local ethics committees (COMETHEA).

### 2.2 Food restriction

Starting after a 4-day habituation period and continuing until the end of the experiment, food was restricted to limit weight gain and to motivate operant behavior. Food (approximatively 20 g/day for male and 15mg/day for females) was given 1 hour after the end of the daily experimental session. Unlimited access to water was provided for the entire duration of experiment. Body weight was 336 ± 1.7 for males and 226 ± 1.7 g for females at the beginning of the experiment and 377 ± 3.2 for males and 249 ± 1.7 g for females at the end of the experiments.

### 2.3 General experimental designs

In this experiment, we aimed to validate the progressive shock strength (PSS) procedure by 1) investigating the effects of motivational manipulations such as hunger (food restriction or satiation), reward size (1 vs 2 pellets of sucrose) and reward quality (sucrose vs chocolate) on punished responding; and 2) directly comparing the effects of these manipulations in our PSS procedure versus a conventional PR procedure. Tests sessions were separated by a minimum of one training session, and each type of test occurred once a week. Sessions were performed 5 days a week and a typical week had the following schedule: Monday: training; Tuesday: PR test; Wednesday: training; Thursday: training and Friday: PSS test. After nine initial operant training sessions, we tested resistance to punishment in 2 PSS sessions and appetitive motivation in 2 PR sessions to establish baseline behavior. The data from these sessions were averaged to provide basal levels of PSS and PR used for statistical analysis. Subsequently, in both PSS and PR, we investigated the effects of 1) hunger (restriction and satiation); 2) quantitative increases in the value of reward (2 pellets vs 1 pellet) and 4) qualitative changes in the value of reward (sucrose vs chocolate pellets). Chocolate pellets (Test Diet 5TUT/1811256) were identical in composition and caloric content to sucrose pellets (Test Diet 5TUT/1811251) but contained chocolate flavor.

For 23h prior to the deprivation and satiation tests, respectively, animals were completely food deprived or had food ad libitum as done previously (Solinas and Goldberg, 2005). To study changes in incentive value of reward, we trained animals with the new condition (2 pellets/reward or 1 chocolate pellet) for 2 days before PSS and PR tests.

24-72h after the last operant sessions, we measured anxiety-related behaviors in an open-field and sensitivity to pain in a hotplate procedure to determine whether individual differences in these processes could have influenced the PSS results.

### 2.4 Food Reinforcement Apparatus and training procedure

Experimental chambers (MedAssociates, www.medassociates.com) were enclosed individually in sound-attenuation chests. Each experimental chamber had a recessed food tray, and two levers in the right wall. The floor was constituted by bars that were connected to shockers that could deliver footshock, with electric current set to 0.45 mA. Each chamber was equipped with a food-pellet dispenser, which could deliver 45 mg pellets to the food tray. Experimental events were controlled by computers using MedAssociates interface and Med-PC IV software; Med-PC code used to conduct the procedures is available upon request. A diode light was present on each lever. One lever was assigned to be the active lever and the corresponding light was used as a conditioned stimulus for food reinforcement. A third diode light was installed on the opposite wall, and its flashing was used as a discriminative stimulus to indicate that food reinforcement would be associated with a foot-shock.

The general training schedule involved 45-min sessions of a fixed-ratio 1 (FR1) schedule of food reinforcement in which one lever press was required to obtain a 45-mg sucrose pellet. Rats initially learned to respond for food during nine sessions under this schedule. During these sessions, food availability was signaled by turning off the house-light, indicating that a single response on the active lever would immediately result in the delivery of one food pellet, accompanied by flashing of the diode light above the lever for 2 sec. Subsequently, the house light was turned on for an additional 18-sec time-out period, during which responding had no programmed consequences. Following the time out, a new trial started and the next response on the right lever was again reinforced. Responses on the inactive lever were recorded but never reinforced.

### 2.5 Self-adjusting progressive shock strength (PSS) procedure

In the PSS procedure, active lever presses resulted in immediate delivery of food rewards and foot-shocks of different strengths by manipulating the shock duration. The PSS consisted of steps in which the shock duration was increased each time the animal obtained 2 separate rewards at a given shock duration. The duration of the first step was 0 sec (no shock), the second step was a low duration of 0.05 sec and subsequent shocks increased of 15% at each step for 20 steps. Thus, the step sequence of durations was: 0, 0.05, 0.06, 0.07, 0.08, 0.09, 0.10, 0.12, 0.13, 0.15, 0.18, 0.20, 0.23, 0.27, 0.31, 0.35, 0.41, 0.47, 0.54, 0.62, 0.71 sec. At the beginning of each shock trial, the light on the side opposite to the levers and food tray was switched on and off intermittently for the entire trial for periods of time proportional to the duration of the shock to signal the presence of shock contingencies according to the formulas: Duration On = 0.1 sec times step number and Duration Off = 2 sec – Duration On. If animals reached the final step, the duration of the shock was not further increased, and all subsequent shock were set at 0.71 sec. If rats did not emit any response for 5 min, shock duration was reset to 0 sec and the shock progression was reinitialized. The 5 min threshold for reset was chosen based on pilot experiments showing that this level allowed most animals to reinitiate their operant behavior within a single session. Thus, animals could at any moment avoid higher strength of shock by limiting the frequency of food reinforcement until shock duration returned to 0 sec, which was signaled by the absence of the blinking light. This feature suppressed but did not completely extinguish operant responding, allowing a rapid return to baseline responding in the following training session. The strength of the shock was determined by the electrical charge in millicoulombs (mC) that an animal would tolerate to obtain food pellets and was calculated by multiplying the fixed current of the shock (0.45 mA) by the duration in sec. The level at which behavior changed abruptly or PSS break point was calculated by the maximal and cumulated strength of the shock received during the session, which, as expected, were highly correlated (r^2^= 0.90 for males and r^2^ = 0.93 for females under basal conditions). We chose the latter parameter for the main analysis because it incorporates both the willingness to receive a given charge unit and the willingness to restart responding after an eventual punishment-induced pause. However, it should be noted that similar results were obtained when the maximal strength of shock obtained was used.

Table 1 summarizes the data obtained in the PSS procedure in terms of responses, duration of shock and total electrical charge received during a session.

**Table 1.**

|  | # Active Responses |  | Total Shock Duration<br>(Sec) |  | Total Electrical<br>Charge (mC) |  |
| --- | --- | --- | --- | --- | --- | --- |
|  | *Males* | *Females* | *Males* | *Females* | *Males* | *Females* |
| **BL** | 120.80 ±<br>1.46 | 97.94±4.024 | - | - | - | - |
| **PSS BL\*** | 45.87±2.94 | 40.50±3.08 | 6.56±0.67 | 5.63±1.37 | 2.95±0.30 | 2.53±0.62 |
| **PSS**<br>**Hungry** | 57.83±2.48 | 59.75±5.88 | 12.79±1.55 | 16.41±4.37 | 5.75±1.04 | 7.38±1.97 |
| **PSS Sated** | 25.33±1.78 | 17.79±2.478 | 1.45±0.23 | 1.38±0.33 | 0.65±0.10 | 0.76±0.02 |
| **PSS 2**<br>**Pellets** | 50.13±2.04 | 32.17±2.38 | 8.01±0.80 | 3.46±0.95 | 3.61±0.36 | 1.56±0.43 |
| **PSS**<br>**Chocolate** | 65.35±2.97 | 55.08±3.80 | 15.49±1.94 | 10.77±2.85 | 6.97±0.87 | 4.84±1.28 |

Mean ± SEM
\* PSS BL was established several times during the experiments. For clarity, here we provide values of the first measure.

### 2.6 Progressive-ratio schedule

Under the progressive-ratio schedule of food reinforcement, the number of responses required to obtain a food pellet increased with each successive food pellet. The steps of the exponential progression were the same as those previously developed by Roberts and colleagues (Richardson and Roberts, 1996) adapted for food reinforcement (Solinas et al., 2003; Solinas and Goldberg, 2005), based on the equation: response ratio = (5e^(0.2^ ^×^ ^reinforcer^ ^number)^) − 5, rounded to the nearest integer. Thus, the values of the steps were 1, 2, 4, 6, 9, 12, 15, 20, 25, 32, 40, 50, 62, 77, 95, 118, 145, 178, 219, 268, 328, 402, 492, 603, and 737. Sessions under the progressive-ratio schedule lasted until 10 min passed without completing a step, which, under basal conditions, occurred within 1 h. This 10 min step-completion threshold produced breakpoints similar to those we observed previously using a 5 min no-response threshold (Solinas et al., 2003; Solinas and Goldberg, 2005).

### 2.7 Anxiety-related behaviors: Open Field

The open-field apparatus (Viewpoint, Lyon, France) consisted of a rectangular arena (50cm wide * 50cm long * 40cm high) of white plexiglass. After a 30-min habituation to the experiment room, rats were placed for 30 min in the arena. Their positions were recorded automatically and in real time by a camera and video tracking software (Viewpoint, Lyon, France). The software defined a virtual square (25cm * 25cm) delimiting the center zone and the border zone. Anxiety was measured as the percentage (%) of time spent in the border (time in the border/time in the border + time in the center * 100) so that more time spent in the border (thigmotaxis) indicated a higher level of anxiety.

### 2.8 Pain sensitivity: Hot plate test

The hot-plate (Ugo Basile, model-DS 37) was maintained at 48 °C (Deuis et al., 2017). After a 10-min habituation to the experiment room, animals were placed into a glass cylinder of 25 cm diameter on the heated surface and 47 cm walls. The latency before escape or jumping, was recorded. Experiments were stopped after a cut-off of 120s to prevent unnecessary pain or tissue damage.

### 2.9 Data and Statistical Analysis

Med-PC data obtained were analyzed using a custom-made, freely available software written in Python, Med_to_csv (https://github.com/hedjour/med_to_csv) that uses raw data files to create complete tables for further analysis in GraphPad Prism. Data were checked for normality distribution on using the Shapiro–Wilk test. PSS data using electrical charge did not show normal distribution and therefore, they were normalized using natural logarithm transformation. For baseline measures of PSS and progressive-ratio break points, we used two-way ANOVA for repeated measures (Factors: sex and procedure) to compare the behavior of male and female rats with Geisser-Greenhouse correction to account for possible violation of sphericity followed by Sidak’s post hoc test. Given that baseline behavior significantly differed between males and females, for subsequent behaviors, we separately compared male and female animals using one-way ANOVA for repeated measures (factor: hunger state) with Geisser-Greenhouse correction followed by Dunnett’s post hoc test or Wilcoxon matched-pairs signed rank test for reward size and quality. Differences were considered significant when p < 0.05.

## 3 RESULTS

### 3.1 PSS and PR break points

The last 3 FR1 training sessions were used to determine a baseline (BL) in the absence of shocks. The number of active responses (and food rewards obtained) was ∼ 121 for male (Fig. A, left) and ∼ 98 for female rats (Fig. 1, right). In the PSS session, the number of pellets decreased to about ∼ 46 in males and ∼ 40 in females, a 62 % and 59 % reduction compared to the BL responding. The PSS break point as measured by the electrical charge sustained was 2.95 ± 0.30 mC in males and 2.53 ± 0.62 mC in females (Table 1). Individual event records (supplementary information, Fig. S1) clearly show that rats titrated their punishment and different rats reached different PSS break points. Indeed, a minority of animals (< 20% under baseline conditions) kept responding for food even when the shock strength was maximal, whereas most (> 80%) of the animals responded until the strength of shock reached a level where responding stopped for at least 5 minutes, leading to a reset of the shock duration to 0 s. Then, most animals started responding again reaching levels that were in a range that was relatively constant for individual animals (average deviation from the median shock strength during baseline < 10%) suggesting that they had a shock strength that was “worth” receiving to obtain their rewards. For PR sessions, the number of steps completed was ∼ 14 in males and ∼ 11 in females and the number of active responses was ∼ 428 in males and ∼ 276 females. Statistical analysis on the number of responses revealed a significant effect of schedule (F(2, 140) =219, p< 0.0001), of sex (F(1,70)= 26.10, p< 0.0001) and a significant schedule * Sex interaction (F(2,140) = 19.41, p < 0.0001). Post-hoc multiple comparison indicated that males and females differed in baseline and PR responding but not in PSS responding (p = 0.31). Because baseline behavior was different in males and females, which can confound the interpretation of results, we decided not to compare them directly in subsequent tests.

**Figure 1.**
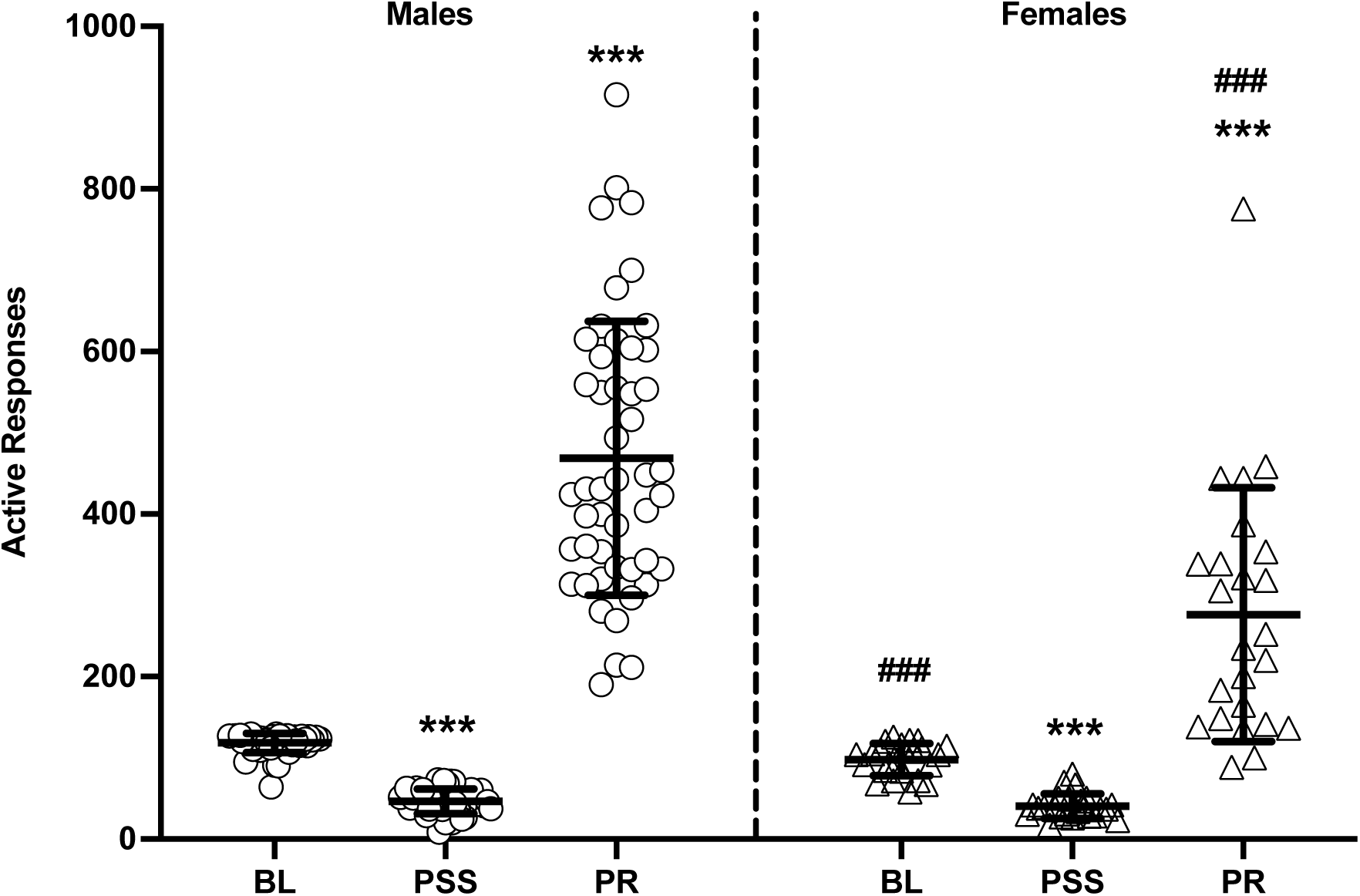
Effects of introducing progressive shock strength (PSS) or progressive ratio (PR) schedules on food-maintained behavior. Number of active responses during baseline (BL), progressive shock strength (PSS) and progressive ratio (PR) sessions in male (left) and female rats (right). Data are expressed as mean ± SD of active responses (Males n = 48; Females = 24). Baseline (BL) was calculated on the last three FR1 training sessions before the first shock session. The values for PSS and PR represent the average of the first 2 separate sessions. Two-way ANOVA for repeated measures followed by Sidak’s post hoc: ***, P < 0.001 PSS and PR vs BL; ###, P < 0.001 males vs females.

Frequency distribution of PSS break points was normally distributed when expressed as natural logarithm of the electrical charge sustained for both males and females with a median value of 0.81 (95% CI: 0.58-1.05) and 0.52 (95% CI: 0.16-0.88), respectively (Fig. 2 A, B). Frequency distribution of PR break points was a conventional Gaussian for males with a median value of 431 (95% CI: 360-549) and a slightly positively skewed Gaussian for females with a median value of 243 (95% CI: 148-339) (Fig. 2 C, D). PSS responding (electrical charge sustained) was not correlated with PR responding (number of responses) in either male or female rats (Fig. 2 E, F) suggesting that these tasks measure somewhat different psychobiological processes.

**Figure 2:**
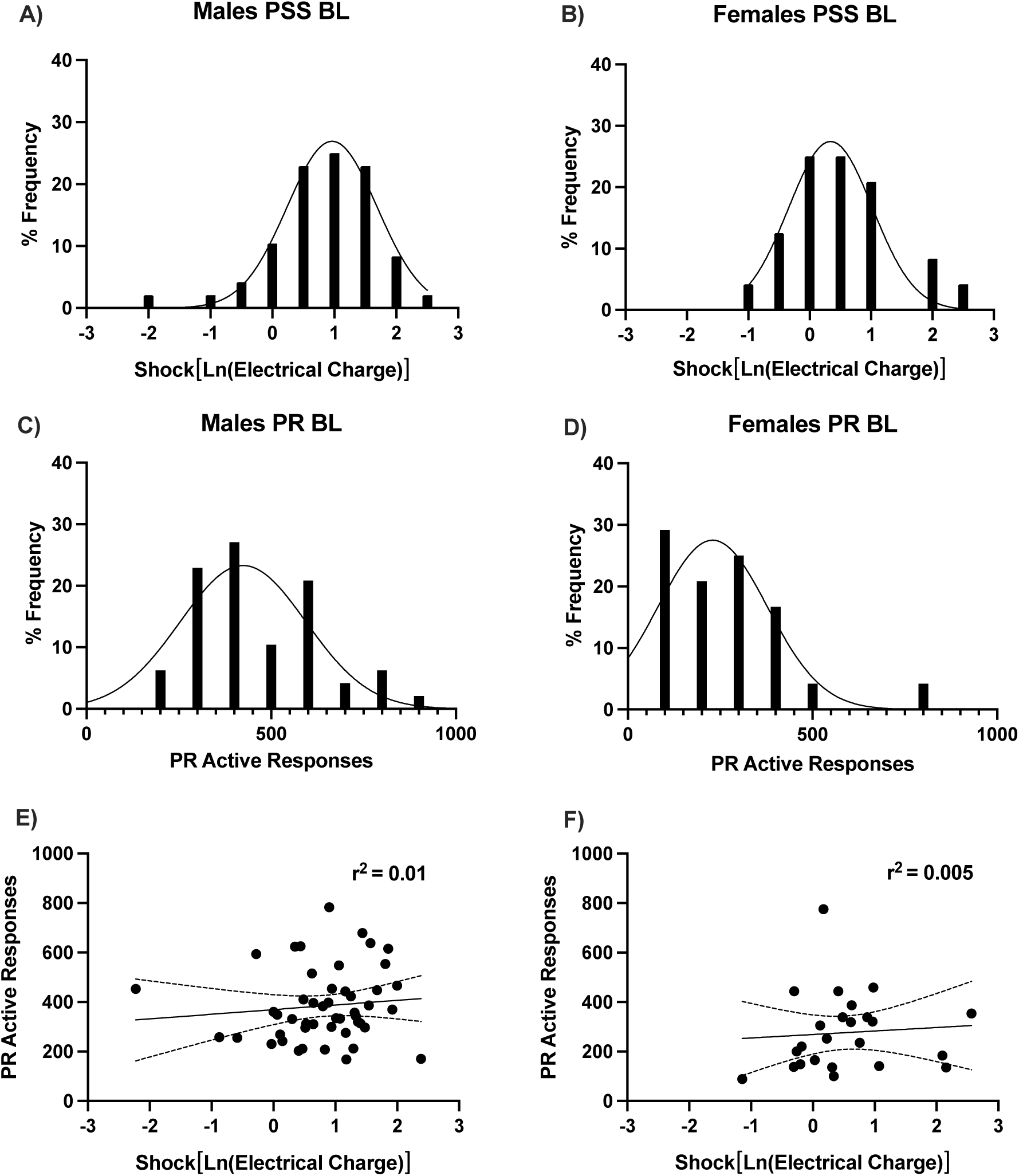
Population distribution for PSS and PR procedures and their correlation. Frequency distribution for behavioral outputs during baseline sessions in the PSS (A, B) and PR (C, D) procedures and their correlation (E, F) in male (left panels: A, C, E) and female rats (right panels: B, D, F). Dotted curves in E and F indicates 95% confidence intervals. The values for PSS and PR represent the average of the first 2 separate sessions.

### 3.2 Effect of hunger on PSS and PR behaviors

To characterize the PSS behavior, we investigated whether it is sensitive to hunger, and we compared the effects of these manipulations on PR. In male rats, hunger induced by food deprivation produced an increase in the PSS break point of 25% in pellets obtained (from ∼ 45 to ∼ 58 pellets) and of 95% in electrical charged received (from 2.95 to 5.75 mC) compared to normal food restriction condition (Fig. 3 A, C, E; Table 1). In contrast, satiation induced by ad libitum pre-session feeding produced a decrease in the PSS break point of 64 % as measured by number of pellets obtained (from ∼ 45 to ∼ 25 pellets) and of 78 % as measured by the electrical charge received (from 2.95 to 0.65 mC) compared to normal food restriction condition (Fig. 3 A, C, E; Table 1). Similarly, in female rats, hunger induced by food deprivation produced an increase in the PSS break point of 50% in pellets obtained (from ∼ 40 to ∼ 60 pellets) and of 250 % in electrical charged received (from 2.53 to 7.38 mC) (Fig. 3B, D, F; Table 1) and food satiation produced a decrease of 56% in the number of pellets obtained (from ∼ 40 to ∼ 18 pellets) and of 75% in the electrical charge (from 2.53 to 0.76 mC) compared to normal food restriction condition (Fig. 3B, D, F; Table 1). Individual event records (supplementary information, Fig. S2) clearly show that, when hungry, rats were willing to receive higher strength of shock than under BL conditions whereas, when sated, they all stop responding at low shock strengths. Statistical analysis revealed a significant effect of hunger state in males (Number of responses: (F(2,47) = 94.11, p< 0.0001; total charge: (F(2,47) = 46.44, p< 0.0001; max charge: (F(2,47) = 112.09, p< 0.0001) and in females (Number of responses: (F(2,23) = 49.15, p< 0.0001; total charge: (F(2,23) = 63.01, p< 0.0001; max charge: ((F(2,23) = 35.15, p< 0.0001).

**Figure 3:**
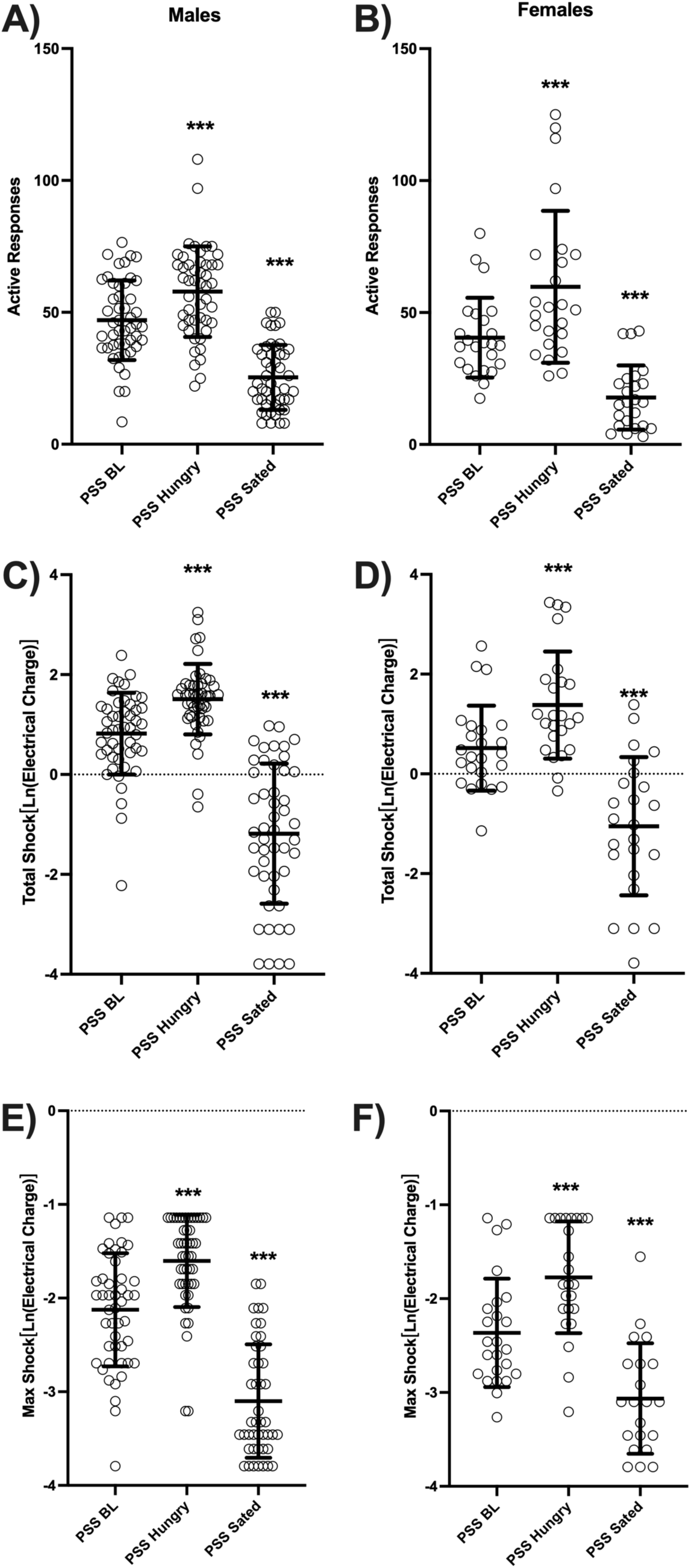
Effects of modulation of hunger on resistance to punishment in the PSS. Resistance to punishment during a PSS session with normal level of food restriction (baseline, PSS BL), after 24h food deprivation (PSS Hungry) or after 24h of access to food ad libitum (PSS Sated) in male (left panels: A, C, E) and female rats (right panels: B, D, F) measured by the number of active responses (A, B), the total electrical charge sustained (C, D) and the max electrical charge sustained (E, F). The values for PSS and PR represent the average of the first 2 separate sessions. One-way ANOVA for repeated measures followed by post hoc Dunnett’s test: ***, P < 0.001 compared to PSS BL.

Consistent with previous reports (Hodos, 1961; Solinas and Goldberg, 2005), PR responding was also sensitive to manipulations of hunger. In males, hunger increased PR responding from ∼ 469 to ∼ 688 responses/session (+47%) and satiation decreased it to ∼ 128 responses/session (−73%) (Fig.4A). In females, hunger increased PR responding from ∼ 277 to ∼ 423 responses/session (+53%) and satiation decreased it to ∼ 64 responses/session (−77%) (Fig. 4B). Statistical analysis of number of responses revealed a significant effect of hunger state in males (F(2,47) = 107.8, p< 0.0001) and in females (F(2,23) = 38.9, p< 0.0001).

**Figure 4:**
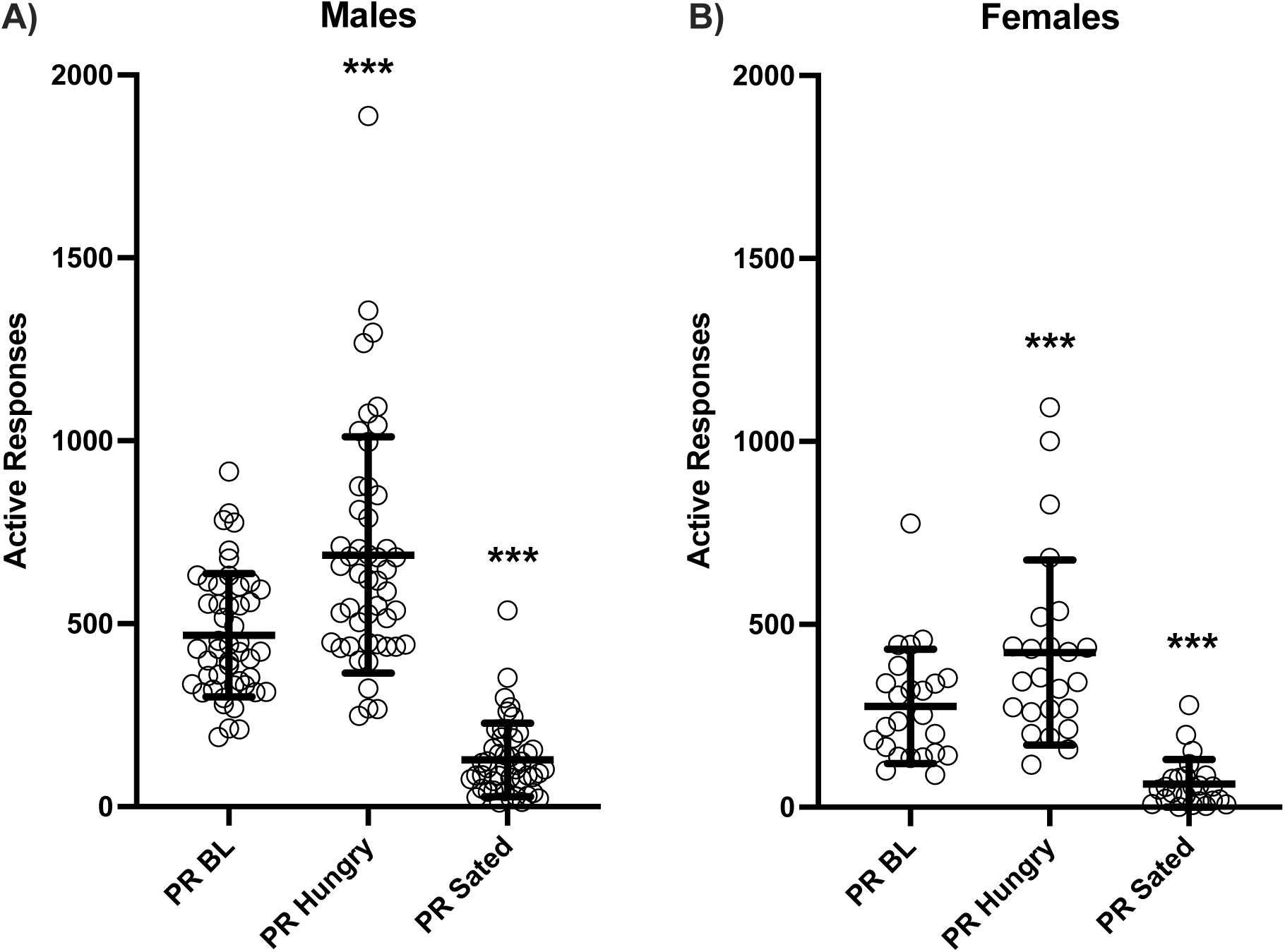
Effects of modulation of hunger on motivation in the PR. Number of active responses during a PR session with normal level of food restriction (baseline, PR BL), after 24h food deprivation (PR Hungry) or after 24h of access to food ad libitum (PR Sated) in male (A) and female rats (B). Data are expressed as mean ± SD (Males n = 48; Females = 24). One-way ANOVA for repeated measures followed by post hoc Dunnett’s test: ***, P < 0.001 compared to PR BL.

### 3.3 Effect of reward size on PSS and PR behaviors

We then investigated whether PSS break points would be sensitive to changes in the reward size. For this, we increased the number of sucrose pellets delivered at each trial to 2 pellets/delivery. Since incentive value needs to be learned, we trained rats for 2 days in this new condition. This manipulation decreased baseline responding because 2 pellets probably induced more rapid satiation. The decrease in baseline was from ∼ 119 to ∼ 104 active responses (and reward deliveries) in males (13%) and from ∼ 98 to ∼ 67 active responses (and reward deliveries) in females (−32%) (data not shown). Compared to this new baseline PSS induced a decrease in both males (from ∼ 104 to ∼ 50 rewards, −50%) and females (from ∼ 67 to ∼ 32, - 52%) (Table 1). PSS break point with 2 pellets did not differ from that with 1 pellet in males (Fig. 5 A, C, E; Table 1; Wilcoxon test: p = 0.14 for active responses, p = 0.47 for total charge and p = 0.11 for max charge) but it was significantly decreased in females (Fig. 4B, D, F; Table 1; Wilcoxon test: p = 0.0053 for active responses, p = 0.025 for total charge and p = 0.16 for max charge).

**Figure 5:**
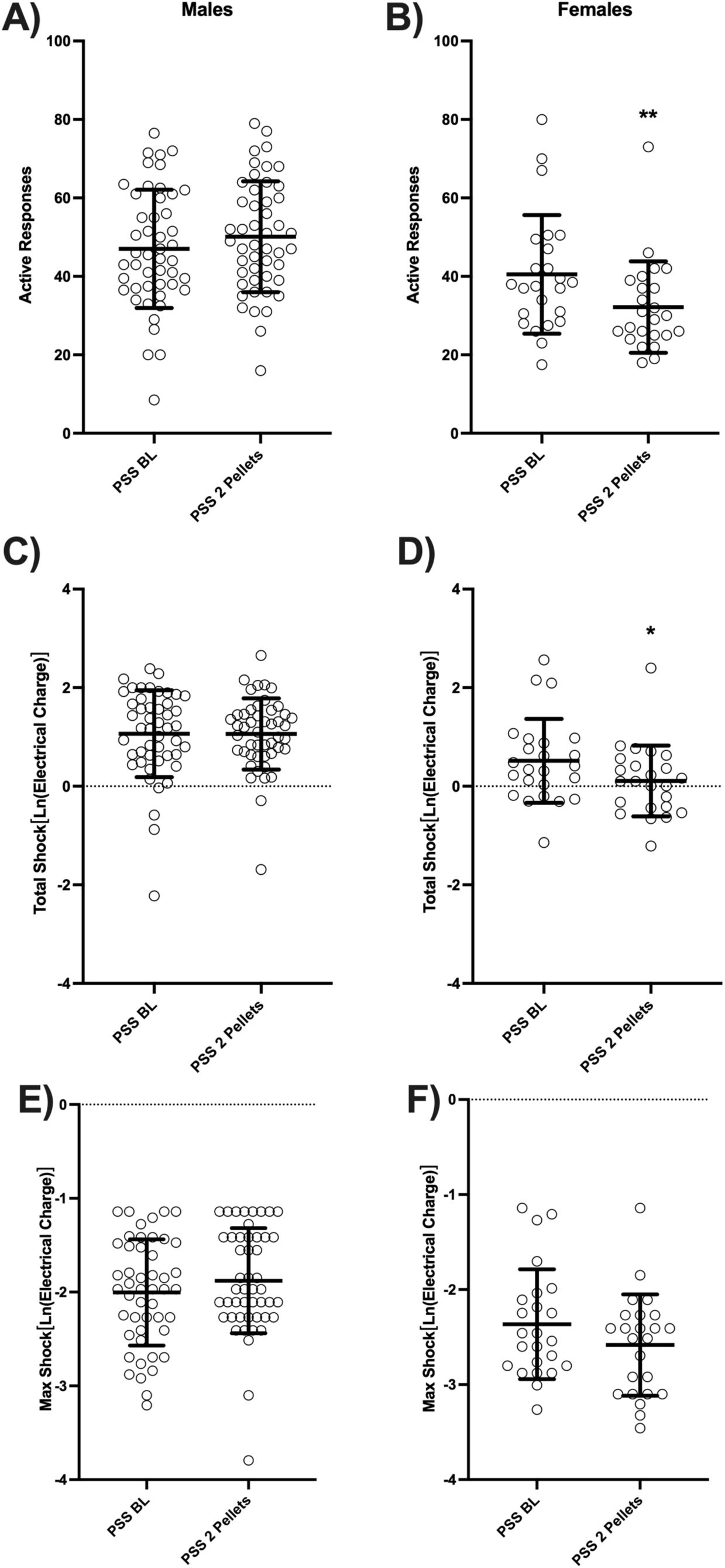
Effects of modulation of reward size on resistance to punishment in the PSS. Resistance to punishment during a PSS session with one pellet/delivery (baseline, PSS BL) or two pellets/delivery (PSS 2 Pellets) in male (left panels: A, C, E) and female rats (right panels: B, D, F) measured by the number of active responses (A, B), the total electrical charge sustained (C, D) and the max electrical charge sustained (E, F). Data are expressed as mean ± SD (Males n = 48; Females = 24). Wilcoxon test: * and **, P < 0.05 and P < 0.01 compared to PSS BL.

PR break point was significantly higher for 2 pellets than for 1 pellet in males (from ∼ 468 to ∼ 734 active responses, +56%; Fig. 6A; Wilcoxon test: p > 0.0001) but was similar in females (∼ 276 vs ∼ 277; Fig. 6B; Wilcoxon test: p = 0.92).

**Figure 6:**
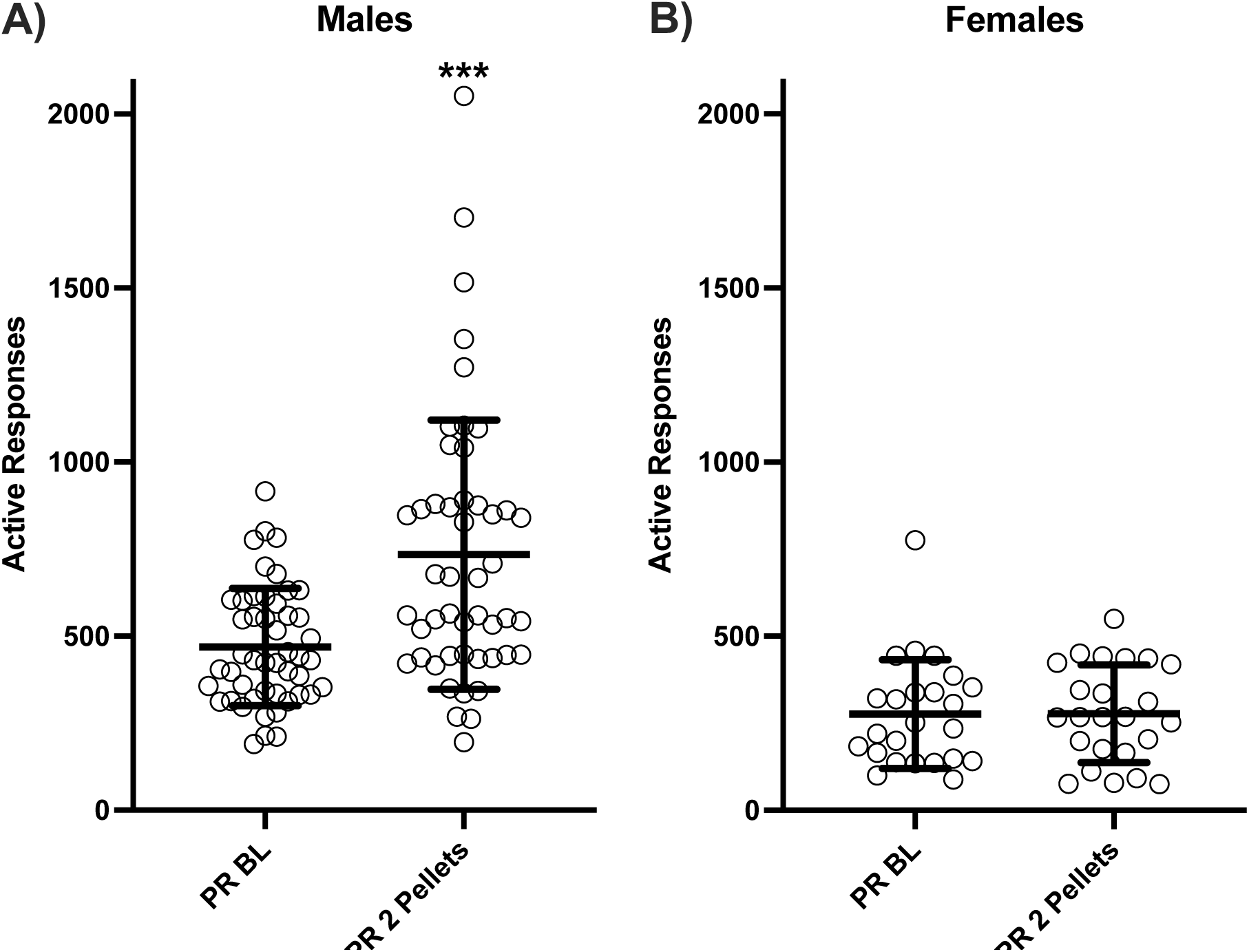
Effects of modulation of reward size on motivation in the PR. Number of active responses during a PR session with one pellet/delivery (baseline, PR BL) or two pellets/delivery (PR 2 Pellets) in male (A) and female rats (B). Data are expressed as mean ± SD (Males n = 48; Females = 24). Wilcoxon test: ***, P < 0.001 compared to PR BL.

### 3.4 Effect of reward quality on PSS and PR behaviors

We then investigated whether PSS break points would be sensitive to changes in the sensory quality of the reward. For this, we allowed rats to self-administer chocolate-flavored pellets. Again, rats were first trained for 2 days in this new condition. This manipulation slightly increased baseline responding in both males and females, from ∼ 119 to ∼ 128 active responses (+7%) in males and from ∼ 98 to ∼ 111 active responses (+13%) in females compared to sucrose pellets (data not shown). Compared to this new baseline PSS induced a decrease in both males (from ∼ 128 to ∼ 65 rewards, −49%) and females (from ∼ 111 to ∼ 55, - 50.5%) (data not shown). More importantly, compared to sucrose pellets, the PSS break point with chocolate pellets was higher in both males (active responses from ∼ 47 to ∼ 65, electrical charge from ∼ 5.92 to ∼ 6.97, Fig. 7 A, C, E; Table 1; Wilcoxon test: p > 0.0001 for active responses and total charge and p = 0.0002 for max charge) and in females (active responses from ∼ 40 to ∼ 55, electrical charge from ∼ 3.40 to ∼ 4.84, Fig. 7B, D, F; Table 1; Wilcoxon test: p > 0.0001 for active responses, p = 0.0004 for total charge and p = 0.0086 for max charge). PR break point was significantly higher for chocolate than for sucrose pellets in both males (from ∼ 468 to ∼ 650 active responses, +39%; Wilcoxon test: p > 0.016) and females (∼ 276 vs ∼535, +94%; Wilcoxon test: p > 0.0001) (Fig. 8A, B).

**Figure 7:**
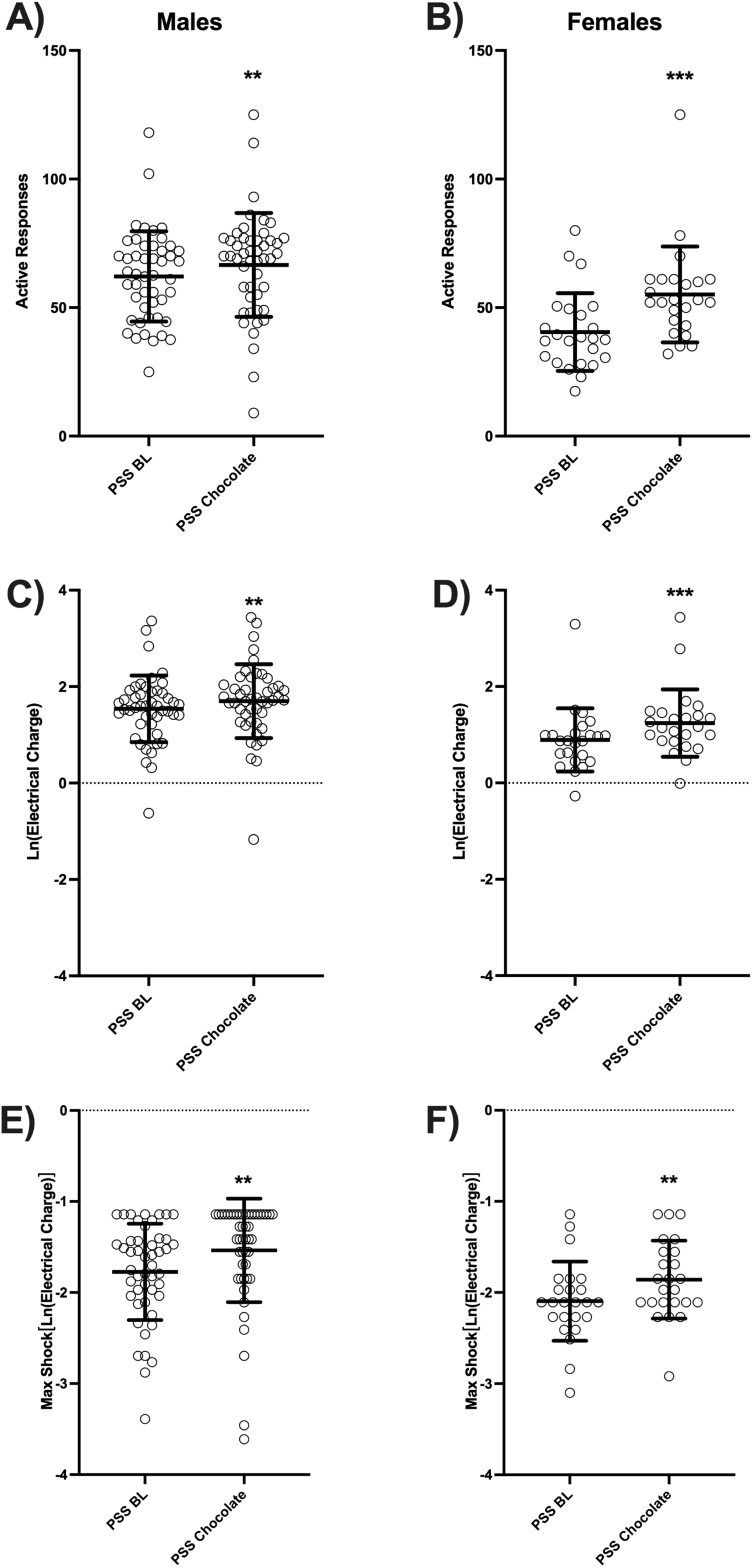
Effects of modulation of reward quality on resistance to punishment in the PSS. Resistance to punishment during a PSS session with delivery of one sucrose (baseline, PSS BL) or one chocolate (PSS Chocolate) in male (left panels: A, C, E) and female rats (right panels: B, D, F) measured by the number of active responses (A, B), the total electrical charge sustained (C, D) and the max electrical charge sustained (E, F). Data are expressed as mean ± SD (Males n = 48; Females = 24). Wilcoxon test:: ** and ***, P < 0.01 and P < 0.001 compared to PSS BL.

**Figure 8:**
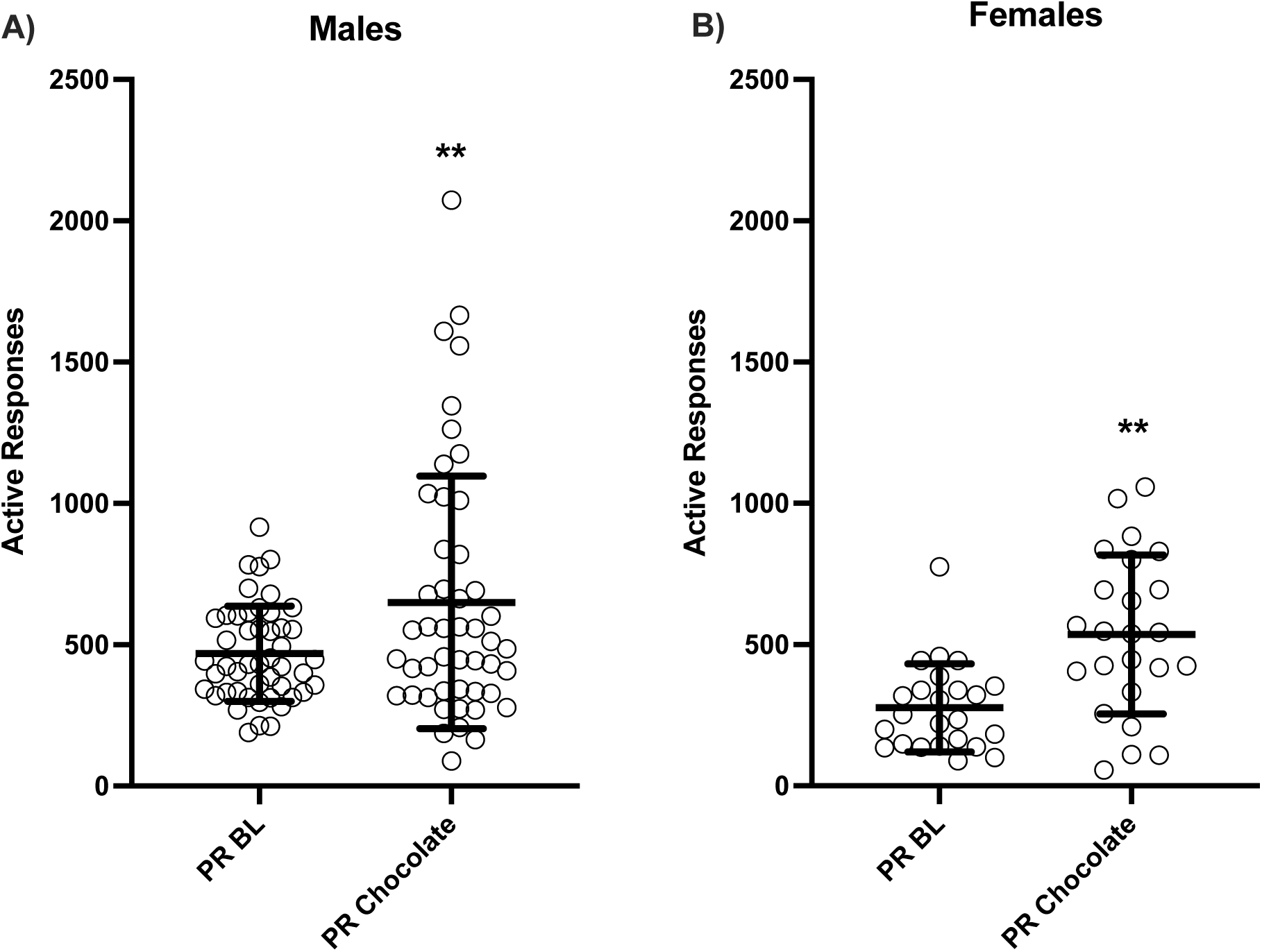
Effects of modulation of reward quality on motivation in the PR. Number of active responses during a PR session with delivery of one sucrose (baseline, PR BL) or one chocolate (PR Chocolate) in male (A) and female rats (B). Data are expressed as mean ± SD (Males n = 48; Females = 24). Wilcoxon test: **, P < 0.01 compared to PR BL.

### 3.5 Anxiety and pain sensitivity

At the end of the operant sessions, we measured anxiety in an open field and pain sensitivity in a hot plate test in order to verify whether these factors could have influenced resistance to punishment. We found that neither anxiety (% time spent in the border) nor pain (latency to escape) was correlated to PSS break points in both males (anxiety r^2^= 0.019 p = 0.34; pain r^2^= 0.003, p = 0.70) and females (anxiety r^2^= 0.048, p = 0.30; pain r^2^= 0.036, p = 0.37) (Fig. S3).

## 4 Discussion

In this study, we developed and characterized a novel self-adjusting progressive shock strength (PSS) procedure to investigate resistance to punishment in which the strength of the punishment is determined by the behavior of the individual animal. Thus, each animal could titrate the amount of shock that it was willing to receive to obtain a food pellet. We show that, in both male and female rats, the PSS break point is sensitive to manipulations of motivation such as hunger and incentive value of the reward. Importantly, interindividual differences in resistance to punishment do not appear to depend on pain sensitivity or anxiety because these measures were not correlated with the PSS break point. Interestingly, although PSS and PR break points were both sensitive to manipulations of motivation, they were not intercorrelated, confirming that these behaviors represent some overlapping but partly distinct constructs and are likely to depend on common but also distinct brain circuits.

In our procedure, shock strength was adjusted for each individual animal in each session, increasing and decreasing depending on the rat’s behavior (increasing in steps while responding was persistent and resetting to 0 when responding ceased for 5 minutes). Most rats took advantage of this feature and resumed food seeking behavior 2 or 3 times in a session (supplementary information, fig. S1). Thus, rats could titrate the levels of shock that they were willing to receive by stopping before they reached a level that was aversive enough to persistently suppress their behavior (Durand et al., 2021). Importantly, our procedure could be considered a form of refinement in the 3Rs (replacement, reduction, refinement) principles of animal research (Hubrecht and Carter, 2019) because animals can control the strength of punishment that they are willing to tolerate. Therefore, the PSS appears particularly fit to stricter ethical standards of animal research that are required by our societies. Finally, the individual calibration of shock levels of the PSS is conceptually similar to what is commonly done in investigate the mechanisms of punishment and compulsivity in humans (Apergis-Schoute et al., 2017; Kanen et al., 2021; Kim and Anderson, 2020) which can increase the translational validity of this procedure.

Shock sessions were repeated but infrequent and sparse. Indeed, we wanted punishment to be rare, similarly to what may happen in humans when negative consequences of drug use or excessive food consumption are rarely (intermittently) experienced after immediate consumption of the reward. In addition, this avoided the cumulative effects of punishment (Durand et al., 2021; Marchant et al., 2018) which can occur when shock sessions are repeated in consecutive sessions. Instead, with sparse shock sessions we were able to rapidly reestablish baseline operant behavior and have several test/retest conditions. Measures of resistance to punishment were highly correlated for the whole experiment (supplementary information, fig. S4) indicating that resistance to punishment is a trait. Therefore, our PSS procedure should allow investigating changes in resistance to punishment longitudinally over long periods of time and in response to behavioral and neurobiological manipulations.

Previous research has shown that rats (and pigeons) will titrate the strength of a constant or pulsing shock to a specific level when the shock strength is contingent on the rate of food-reinforced responding (Rachlin, 1972; Rachlin and Loveland, 1971); this level was stable within individuals but differed across individuals. More recent studies investigating footshock punishment found bimodal population distributions with procedures using a fixed shock strength (Belin et al., 2011; Domi et al., 2021; Jean-Richard-Dit-Bressel et al., 2019; Marchant et al., 2018) and continuous Gaussian distribution of responding with procedures using increasing levels of punishment with food (Datta et al., 2018), cocaine (Datta et al., 2018) and alcohol (Marchant et al., 2018) as reinforcers. The distribution in our self-adjusting PSS procedure was a positively skewed Gaussian when the electrical charge was plotted (data not shown) but it was a normal Gaussian when the strength was expressed as the natural logarithm of the electrical charge. This is consistent with Stevens’ power law that states that sensation strength and stimulus intensity are described by a power function (Stevens, 1957) and with previous studies investigating the consequences of different shock strengths (Leander, 1973). This suggests that when animals can self-titrate the level of punishment, individual levels of shock resistance are log-normally distributed whereas bimodal distribution of shock resistance may at least in part result from imposing a fixed strength that artificially divide the population in two groups of resistant or sensitive individuals.

Hunger is a potent motivation of food seeking and taking behavior because of the intrinsic evolutionary value of 1) working to obtain food when energy stocks are low and 2) avoiding excessive risk or unnecessary exertion when energy stocks are already sufficient. In this study, both PSS and PR break points were very sensitive to hunger manipulation, and rats dramatically increased their break points when hungry and decreased it when sated. This is consistent with previous findings in rats with PR (Hodos, 1961; Solinas and Goldberg, 2005) and with punishment (Jacobs and Moghaddam, 2020; Storms, 1963). Incentive value of food is also an important motivator of food seeking and taking. In different versions of choice experiments, rats rapidly learn to perform actions that lead to the delivery of higher quantity of reward and this preference can be used to investigate the effects of manipulations of delay, risks, etc (Evenden and Ryan, 1996; Floresco et al., 2008; Wahab et al., 2018), and higher concentrations of a liquid reward maintain higher break points in the PR (Hodos, 1961; Reilly, 1999). In our procedure, manipulations where reward size was increased produced inconsistent results. For PSS, no effect was found in males but a decrease in break point was found in females whereas for PR, an increase was found in males and no effect was found in females. These results are likely to be highly influenced by satiation effects associated with doubling the reward size; these effects were stronger in females compared to males because of metabolic needs and were stronger in the PSS compared to PR because in this procedure rats obtain higher number of rewards in the PSS (10-20 deliveries/session in the PR and 30-50 for PSS). For example, it has been shown that PR responding becomes independent from concentration of liquid rewards when rats are water deprived (Reilly, 1999). Therefore, we cannot interpret these results purely in terms of incentive value. In contrast to reward size, manipulations of reward quality (giving chocolate pellets instead of sucrose) led to clear increases in both PSS and PR break points in males and females. Thus, both PSS and PR are sensitive to changes in incentive value of the reward if satiation levels are maintained at a constant level.

Differences in sensitivity to punishment may be explained by different behavioral mechanisms such as differences in reward/motivation processes, sensitivity to the aversive stimulus, pavlovian learning, encoding the relation between response and the punishment or the inability to control behavior (Jean-Richard-Dit-Bressel et al., 2019). In an elegant study using a conditioned punishment task that distinguished between different aspects of punishment insensitivity, Jean-Richard-Dit-Bressel et al. (2019) found that insensitivity to punishment is not related to reward/motivation or Pavlovian fear. The authors argue that individual differences in resistance to punishment is related to deficits in encoding the contingency between behavior and the delivery of shock and not to differences in behavioral inhibition because punishment-resistant rats showed normal suppression of behavior by Pavlovian fear. Although our study was not designed to investigate the specific mechanisms of punishment, the fact that punishment and motivation breakpoints were not intercorrelated under basal conditions (fig. 2) and throughout the experiment (supplementary information, fig. S4), is in agreement with the interpretation that resistance to punishment is not simply due to reward/motivation processes. These results are also in agreement with previous papers that directly investigated both measures for drugs such as cocaine (Datta et al., 2018; Pelloux et al., 2007; but see Venniro et al., 2018), food rewards (Datta et al., 2018) or direct stimulation of VTA dopamine neurons (Pascoli et al., 2015). It is possible that motivation in progressive ratio schedules is mostly dependent on reward neurotransmitters such as dopamine (Cagniard et al., 2006; Ko and Wanat, 2016; Kravitz et al., 2012; Randall et al., 2012), endoopioids and endocannabinoids (Solinas and Goldberg, 2005) whereas resistance to punishment depends also on brain serotonin levels (Cohen et al., 2015; Pelloux et al., 2012); it is also possible that the balance between these two systems (Palminteri and Pessiglione, 2017) and/or the involvement of other neurotransmitters such as glutamate and GABA (Stephenson-Jones et al., 2020) participate in determining individual differences. Importantly, consistent with previous studies (Degoulet et al., 2021; Li et al., 2021), sensitivity to aversion does not seem to be responsible for differences in punishment because pain sensitivity in the hot-plate test did not differ between resistant and sensitive rats. Future studies are needed to investigate the precise mechanisms responsible for resistance to punishment in the PSS procedure.

Differences exist between males and females in the incidence and prevalence of psychiatric disorders and namely drug addiction and eating disorders (Earls, 1987; O’Brien and Anthony, 2005; Solmi et al., 2021) and have also been reported in animal models (Asarian and Geary, 2013; Becker and Koob, 2016). Previous studies investigating sex differences in the motivation for food have reported no difference in PR responding (van Hest et al., 1988) or slightly higher responding in males compared to females that could be explained by differences in the procedure, baseline responding and body weight (Datta et al., 2017). On the other hand, sex differences have been reported in punished-guided actions with females being more sensitive than males when punishment was probabilistic and less sensitive when it was certain (Chowdhury et al., 2019). In this study, we found differences in baseline behaviors, with males having higher food intake and PR responding, consistent with males having higher body weight, vigor, and metabolic needs. Because of these differences, which could confound the interpretation of the results, we avoided direct comparison between males and females and instead we analyzed their behavior separately. Importantly, we found that manipulations of hunger and incentive value of food produced qualitatively similar effects on PSS and PR in males and females.

Although our PSS procedure appears to have several strengths, it may have some limitations compared to other procedures. For example, several punishment procedures use probabilistic punishment schedules in which only a percentage (often 50%) of responses are punished (Jacobs and Moghaddam, 2020; Marchant et al., 2018; Pascoli et al., 2015). This has the advantage of modeling the fact that punishment in humans is uncertain and, in addition, it avoids animals to completely stop responding in punishment sessions. In our procedure, adding a probability factor would have reduced the number of times a rat could experience a given strength of shock or it would have required more trials at a given strength before changing it, which would have undoubtedly complicated the procedure and the interpretation of the results. Another issue that is addressed directly by other procedures but not ours, is the distinction between seeking and taking (Jacobs and Moghaddam, 2020; Pascoli et al., 2015; Pelloux et al., 2007). With food reinforcement, after pressing the “taking” lever, rats have to perform a further action to consume the pellet, therefore, it could be argued that pressing the lever in our procedure is reward seeking whereas eating the pellet is reward taking. It could be interesting to investigate whether similar or different results would be obtained in the PSS procedure if a seeking lever were added, and punishment occurred on this lever well before having access to the food. In general, whereas the parameters (values of shock, progression, number of repetitions, etc) we chose for our PSS procedure provided a sensitive measure of resistance to punishment of responding for food reward, it remains to be investigated whether other parameters would provide more sensitive measures of resistance to punishment of responding for food reward or for other natural, pharmacological or artificial rewards.

In conclusion, we developed and characterized a novel operant self-adjusting procedure to investigate resistance to punishment in rats. This procedure provides in a single session an individual value, the PSS break point, that reflects the willingness of the animal to seek and take a reward despite negative consequences, a characteristic of compulsive behaviors and addiction. The PSS procedure may be a useful, efficient tool to investigate neurobiological mechanisms of punishment and it could also be easily adapted to the investigation of compulsive seeking in drug addiction processes.

## Acknowledgements

We thank Youna Vandaele and Pauline Belujon for helpful comments on a previous version of the manuscript. This study has benefited from the facilities and expertise of PREBIOS platform (Université de Poitiers).

## Author contributions

SD: Analysis of data, revising the article; VL, Acquisition and analysis of data, revising the article; JEL and MH, Analysis, preparation of the figures and revising the article; LVP, Conception and programming of the procedure, interpretation of data, revising the article; NT: Conception, revising the article; MS: Conception and design, Analysis and interpretation of data, Drafting and revising the article

