## Supplementary information for "A self-adjusting, progressive shock strength procedure to investigate resistance to punishment: characterization in male and female rats"

**SUPPLEMENTARY INFORMATION**
**Supplementary figures and legends**

**Fig. S1. Individual event records for animals from the 3 cohorts (2 X 24 males and 24 females) in the PSS procedure under baseline conditions.** Each page represents a cohort of rats: 1) 24 males; 2) 24 males and 3) 24 females. Data from the second session of baseline PSS are plotted. Each panel represents the behavior of an individual rat in a 45 min session. The X axis shows the time within the session, and the Y axis shows the intensity of each subsequent shock received. Circles show the delivery of the reward and the shock. Note that if rats stopped responding for 5 min, the shock level was reset to 0. The shock condition was signaled by a flashing light whose blinking speed was proportional to the intensity of the shock.

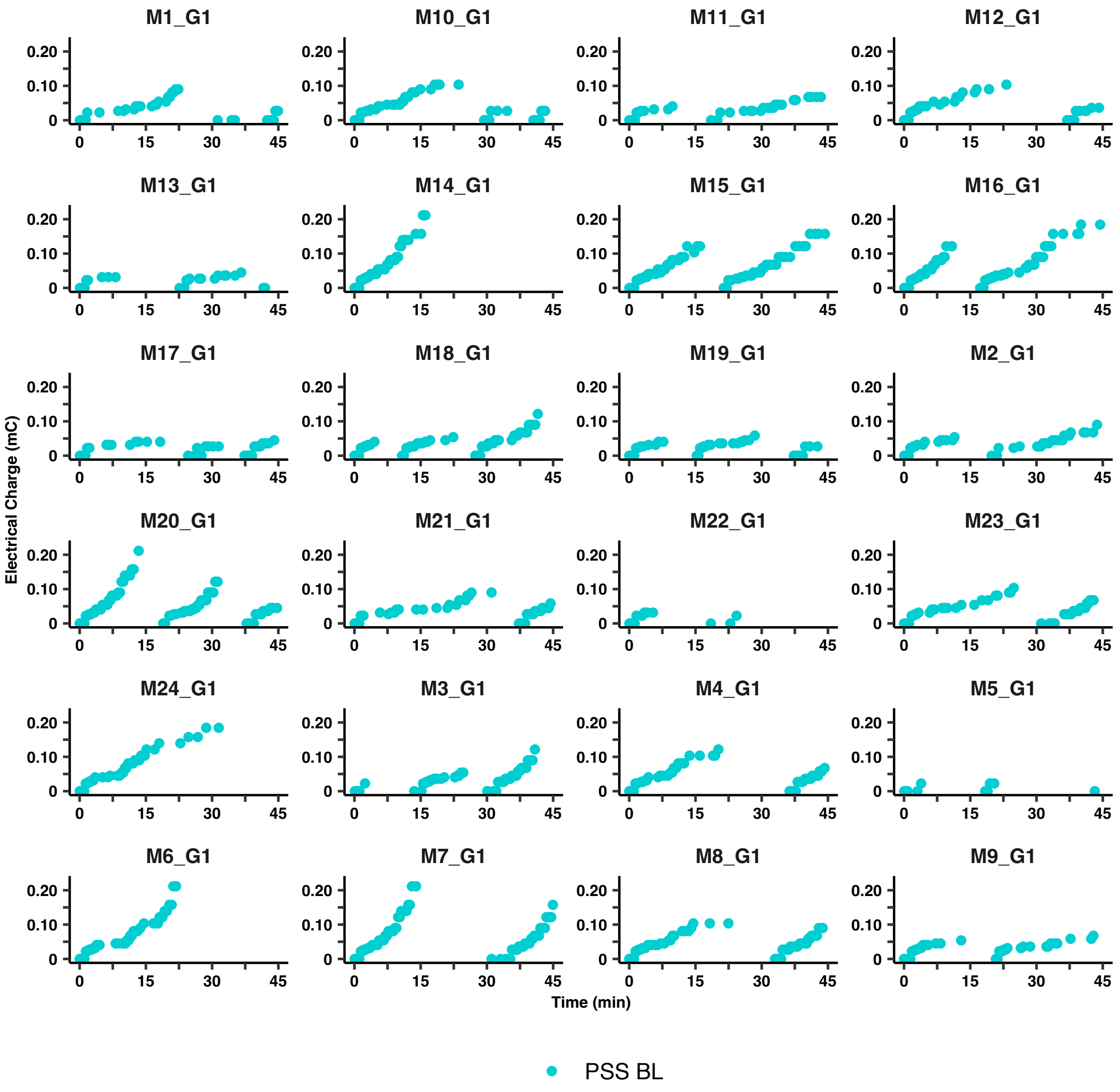

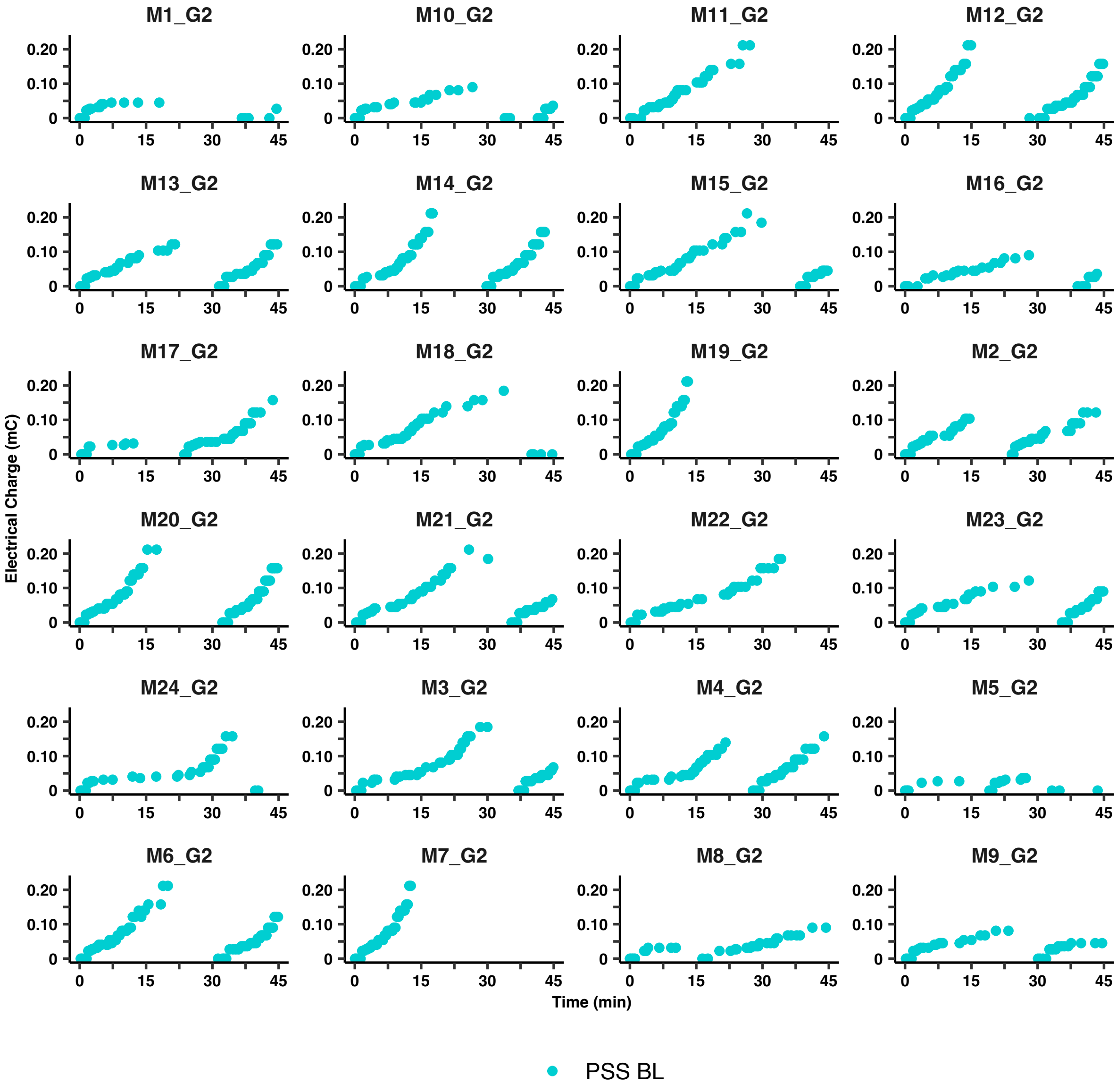
M1_G2
M10_G2
M11_G2
M12_G2
M13_G2
M14_G2
M15_G2
M16_G2
M17_G2
M18_G2
M19_G2
M2_G2
M20_G2
M21_G2
M22_G2
M23_G2
M24_G2
M3_G2
M4_G2
M5_G2
M6_G2
M7_G2
M8_G2
M9_G2
Electrical Charge (mC)
Time (min)
0.20
0.10
0
0
15
30
45
PSS BL

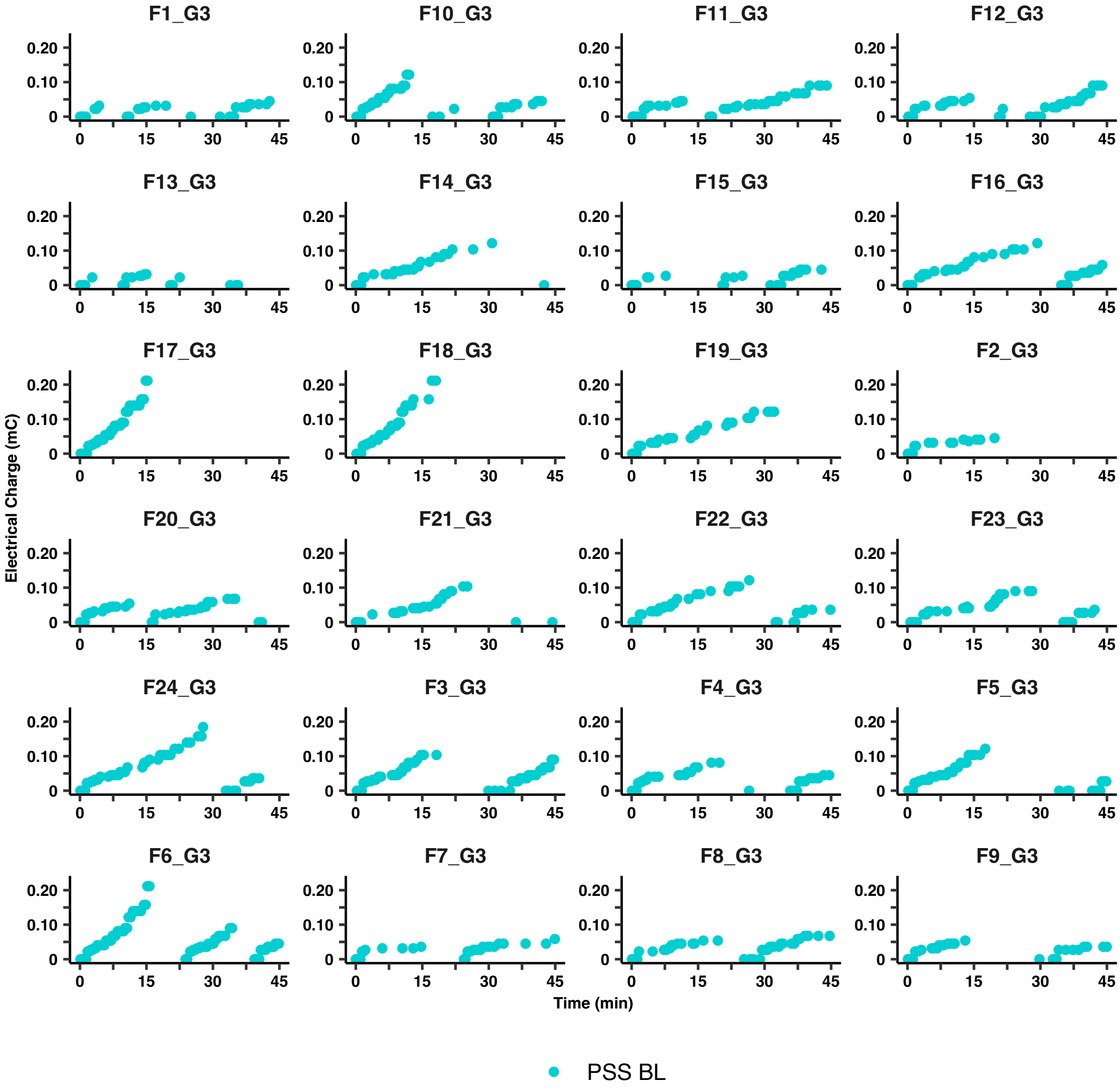
F1_G3
F10_G3
F11_G3
F12_G3
F13_G3
F14_G3
F15_G3
F16_G3
F17_G3
F18_G3
F19_G3
F2_G3
F20_G3
F21_G3
F22_G3
F23_G3
F24_G3
F3_G3
F4_G3
F5_G3
F6_G3
F7_G3
F8_G3
F9_G3
Electrical Charge (mC)
Time (min)
0.20
0.10
0
0
15
30
45
PSS BL

**Fig. S2. Individual event records for animals from the 3 cohorts in the PSS procedure under baseline, restriction and satiation conditions.** Each page represents a cohort of rats: 1) 24 males; 2) 24 males and 3) 24 females. Each panel represents the behavior of an individual rat in a 45 min session. The X axis show the time within the session and the Y axis shows the intensity each subsequent shock received. Circles show the delivery of the reward and the shock. Note that if rats stopped responding for 5 min, the shock level was reset to 0. The shock condition was signaled by a flashing light whose blinking speed was proportional to the intensity of the shock. Baseline data are the same as shown in fig S1.

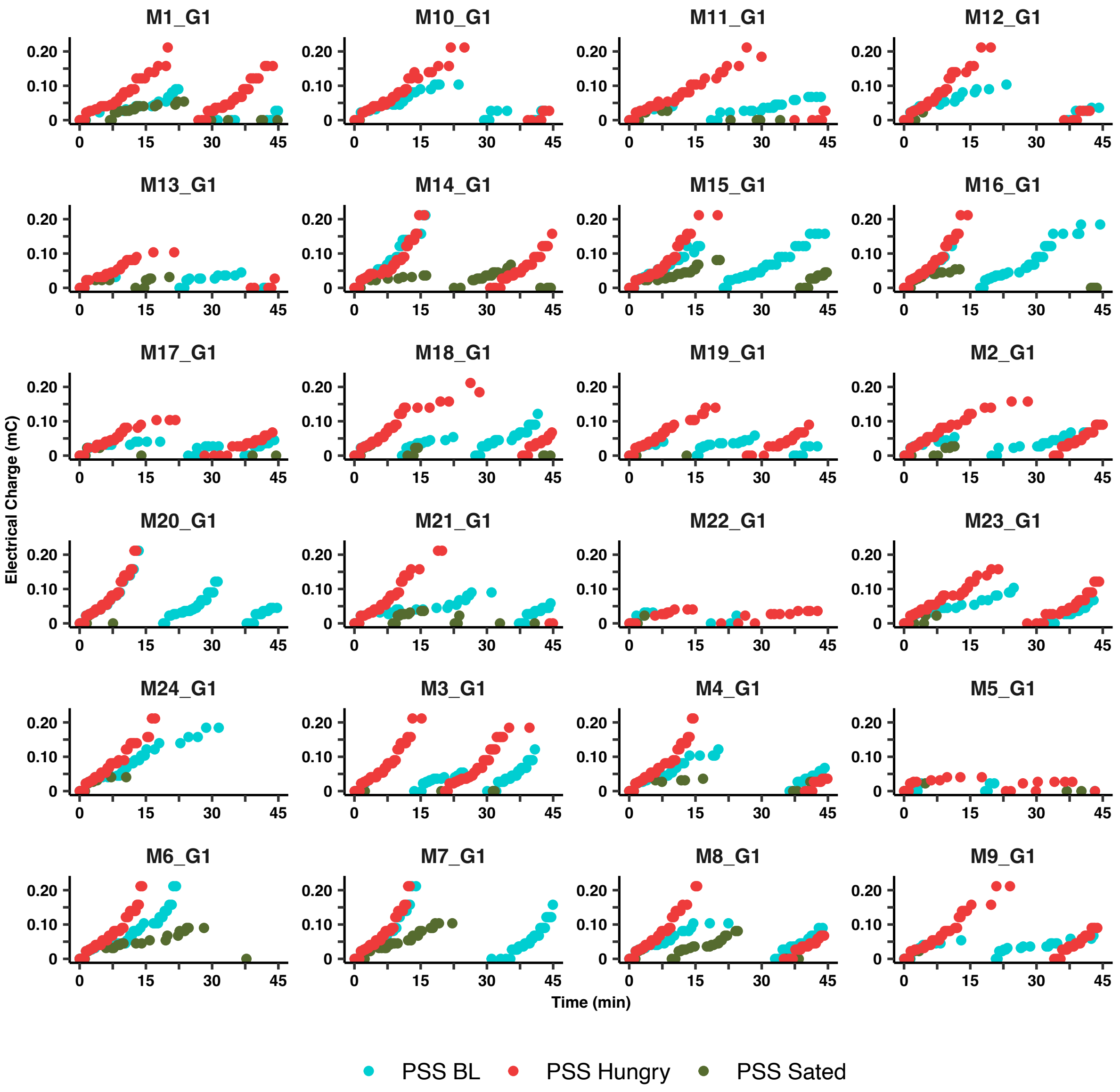

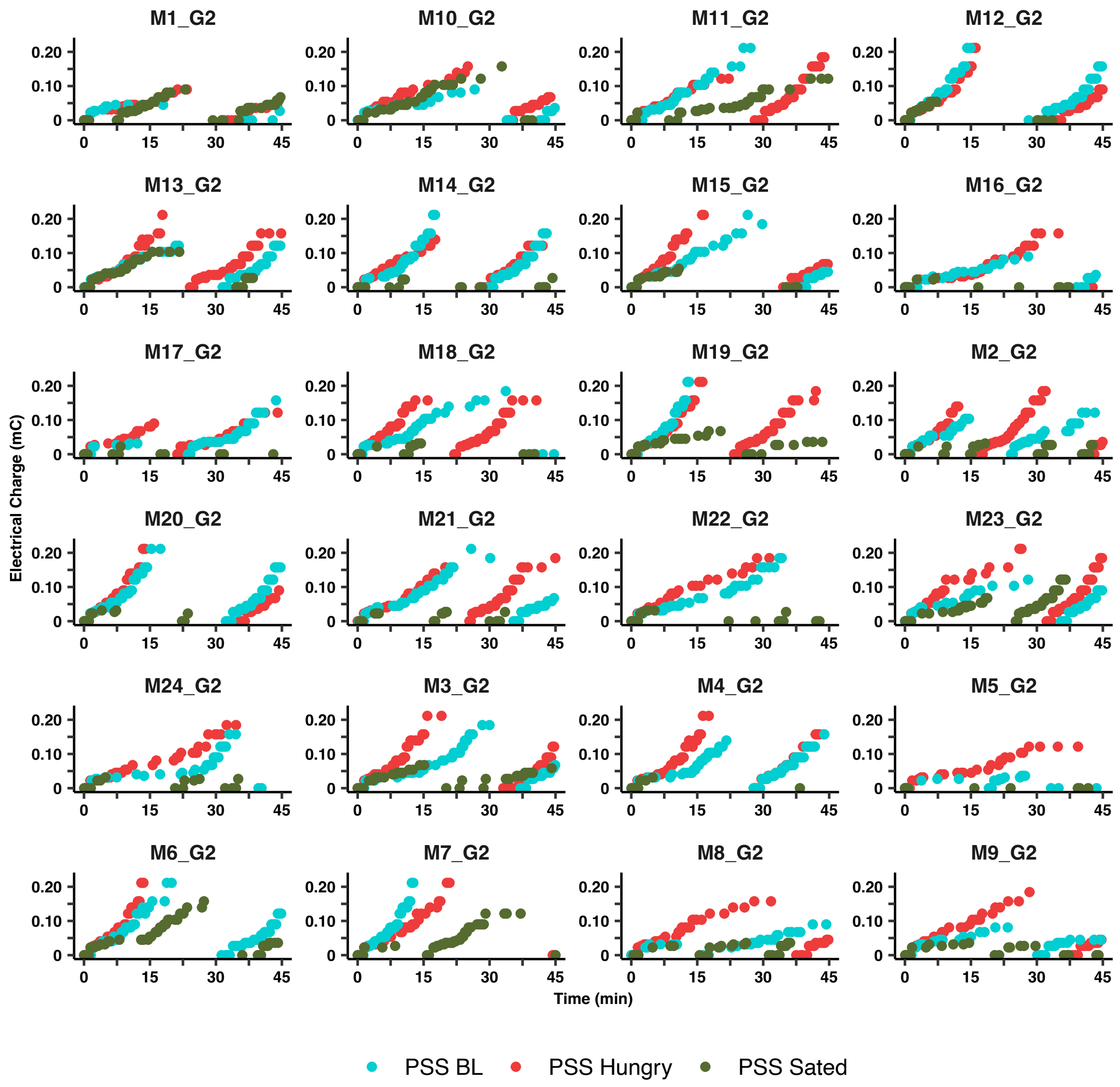
M1_G2
M10_G2
M11_G2
M12_G2
M13_G2
M14_G2
M15_G2
M16_G2
M17_G2
M18_G2
M19_G2
M2_G2
M20_G2
M21_G2
M22_G2
M23_G2
M24_G2
M3_G2
M4_G2
M5_G2
M6_G2
M7_G2
M8_G2
M9_G2
Electrical Charge (mC)
0.20
0.10
0
0
15
30
45
Time (min)
PSS BL
PSS Hungry
PSS Sated

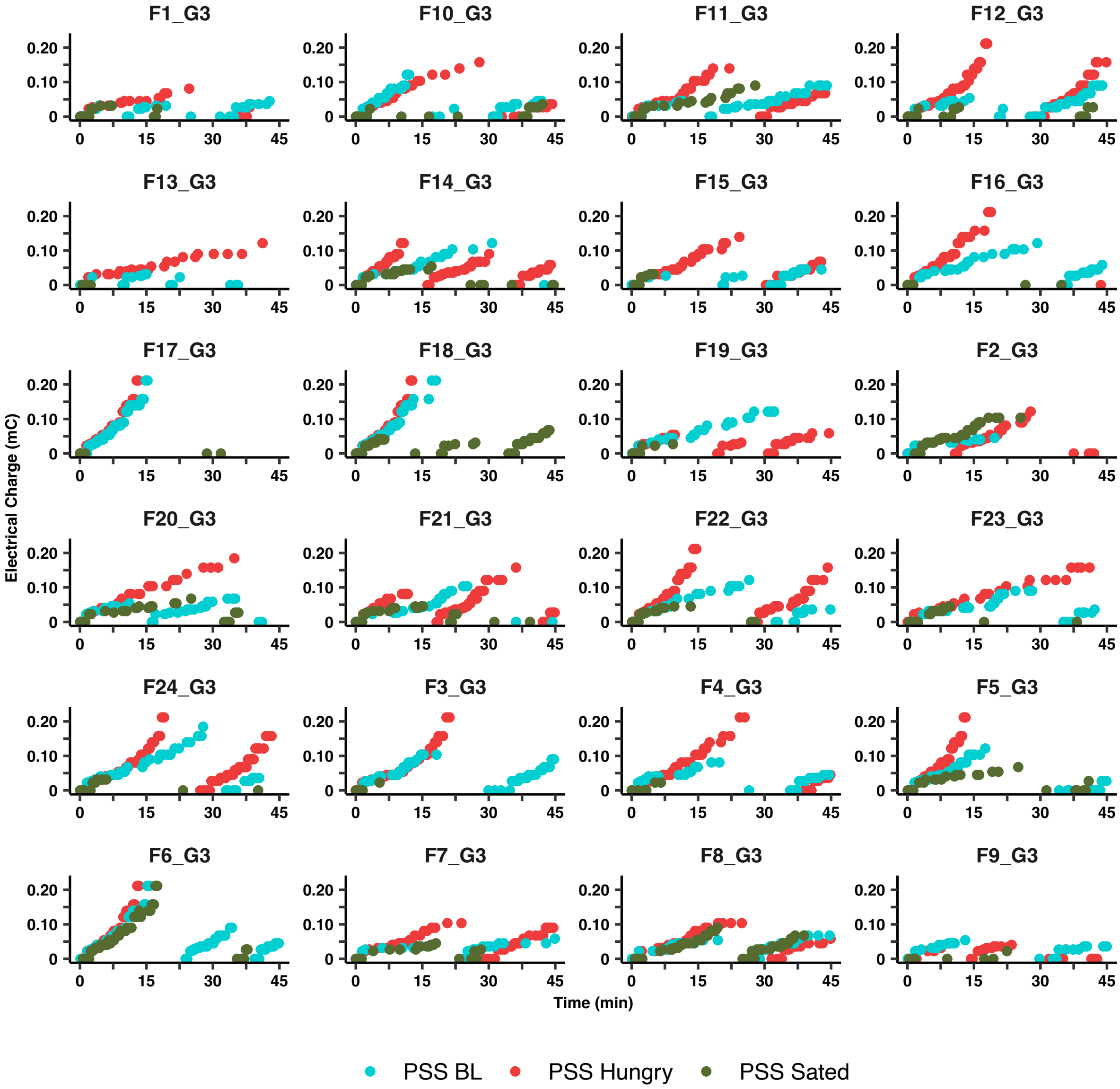
F1_G3
F10_G3
F11_G3
F12_G3
F13_G3
F14_G3
F15_G3
F16_G3
F17_G3
F18_G3
F19_G3
F2_G3
F20_G3
F21_G3
F22_G3
F23_G3
F24_G3
F3_G3
F4_G3
F5_G3
F6_G3
F7_G3
F8_G3
F9_G3
Electrical Charge (mC)
Time (min)
PSS BL
PSS Hungry
PSS Sated

**Fig. S3. Lack of correlation between anxiety levels and pain levels and PSS break points.** Linear regression between anxiety levels measured as % time spent in the center of an open field and PSS break points in A) male and B) female rats. Linear correlation between pain sensitivity measured as latence to escape from a hot-plate and PSS break points in C) male and D) female rats. Dotted curves in E and F indicates 95% confidence intervals.

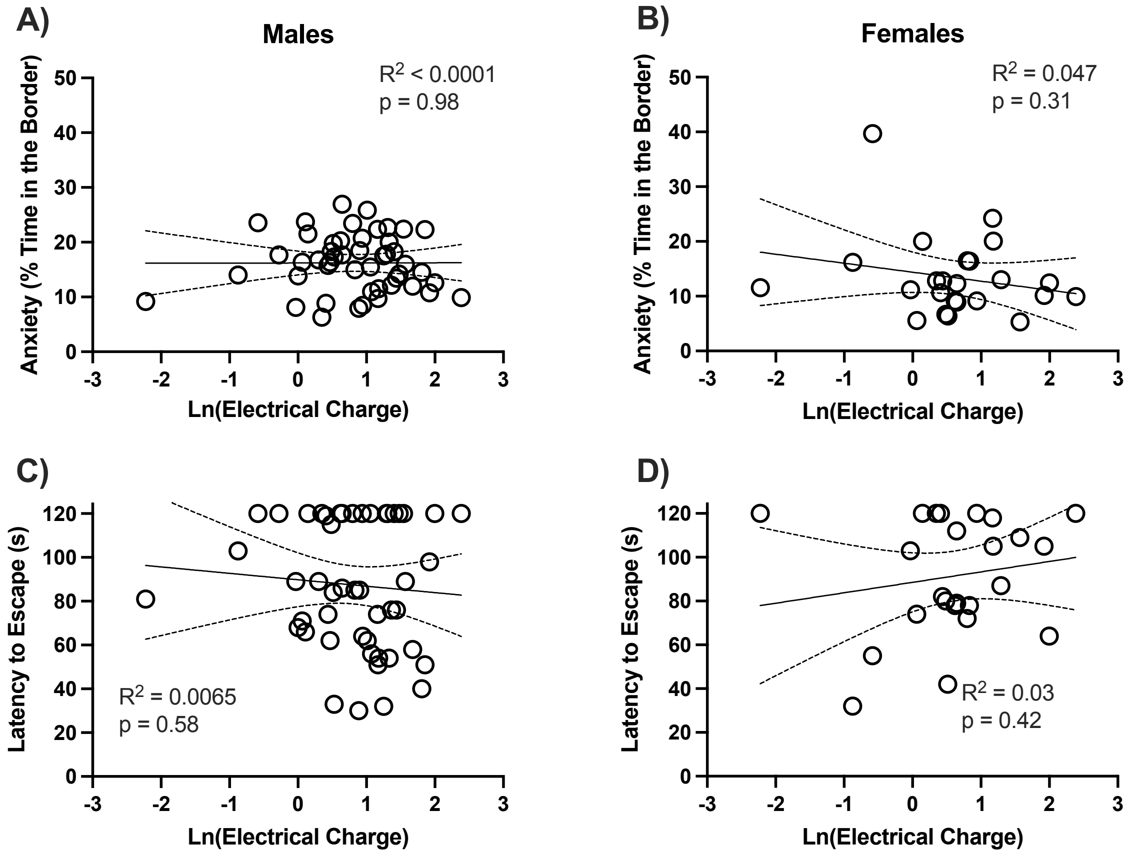

**Fig. S4. Heatmap of within-subject correlation of PSS and PR values under different conditions in Male and Female rats.** Notice that PSS break points were correlated throughout the experiments that lasted several months, showing consistency over time within individual rats. Similarly, PR break points were correlated over time. However, PSS and PR levels show no or little correlation under the different conditions tested.

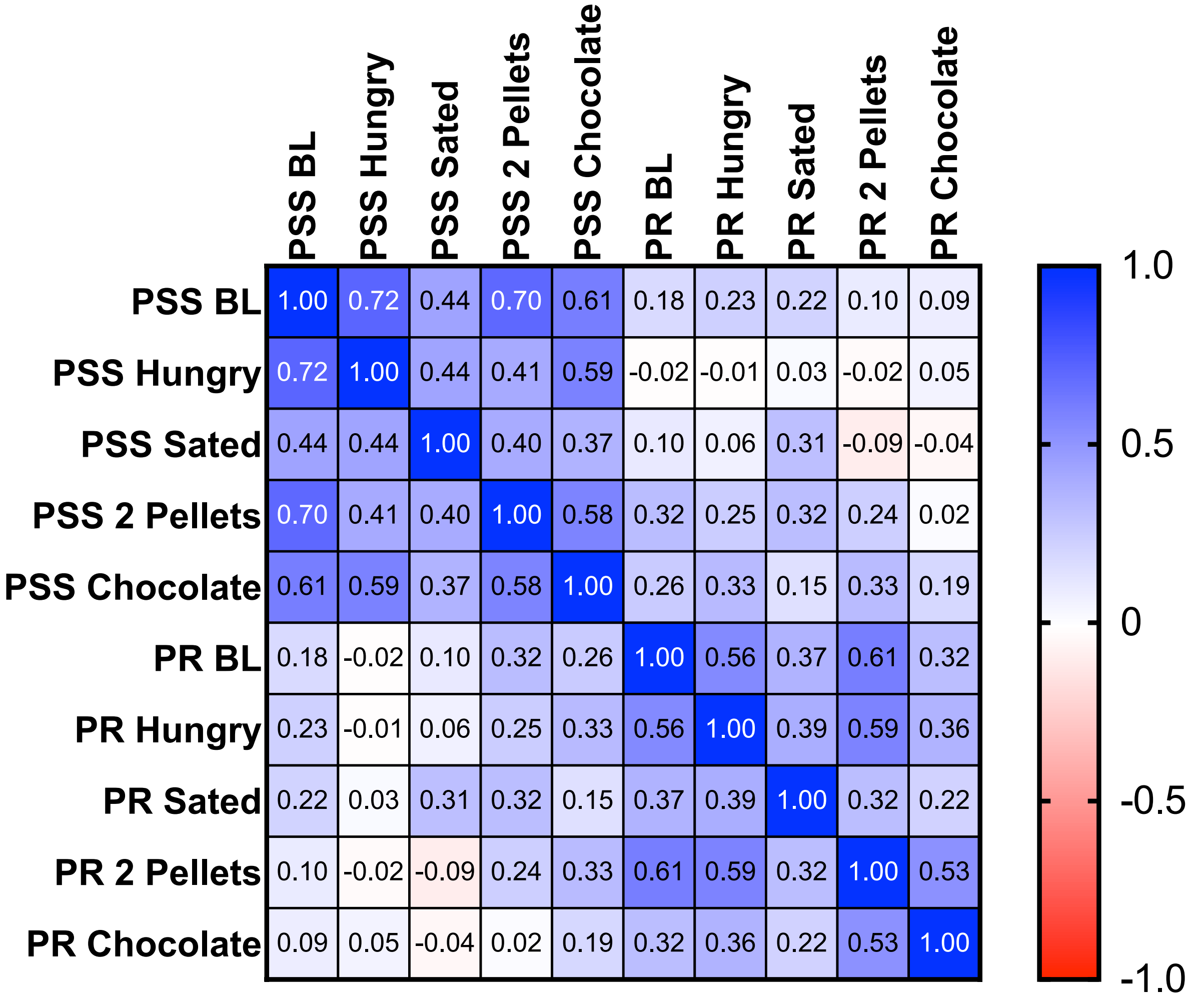
